# Alternate routes to acetate tolerance lead to varied isoprenol production from mixed carbon sources in *Pseudomonas putida*

**DOI:** 10.1101/2024.10.29.620962

**Authors:** Guilherme M. V. de Siqueira, Aparajitha Srinivasan, Yan Chen, Jennifer W. Gin, Christopher J. Petzold, Taek Soon Lee, María-Eugenia Guazzaroni, Thomas Eng, Aindrila Mukhopadhyay

## Abstract

Lignocellulose is a renewable resource for the production of a diverse array of platform chemicals, including the biofuel isoprenol. While this carbon stream provides a rich source of sugars, other organic compounds, such as acetate, can be used by microbial hosts. Here we examined growth and isoprenol production in a *Pseudomonas putida* strain pre-tolerized (“PT”) background where its native isoprenol catabolism pathway is deleted, using glucose and acetate as carbon source. We found that PT displays impaired growth in minimal medium containing acetate, and often fails to grow in glucose-acetate medium. Using a mutant recovery-based approach, we generated tolerized strains that overcame these limitations, achieving fast growth and isoprenol production in the mixed carbon feed. Changes in the glucose and acetate assimilation routes, including an upregulation in PP_0154 (SpcC, succinyl-CoA:acetate CoA-transferase) and differential expression of the gluconate assimilation pathways, were key for higher isoprenol titers in the tolerized strains, while a different set of mechanisms were likely enabling tolerance phenotypes in media containing acetate. Among these, a coproporphyrinogen-III oxidase (HemN) was upregulated across all tolerized strains, and in one isolate required for acetate tolerance. Utilizing a defined glucose and acetate mixture ratio reflective of lignocellulosic feedstocks for isoprenol production in *P. putida* allowed us to obtain insights into the dynamics and challenges unique to dual carbon source utilization that are obscured when studied separately. Together, this enabled development of a *P. putida* bioconversion chassis able to use a more complex carbon stream to produce isoprenol.

**Importance:** Acetate is a relatively abundant component of many lignocellulosic carbon streams and has the potential of being used together with sugars, especially in microbes with versatile catabolism such as *P. putida*. However, the use of mixed carbon streams necessitates additional optimization. Further, the use of *P. putida* for the production of the biofuel target, isoprenol, requires the use of engineered strains that have additional growth and production constraints when cultivated in acetate and glucose mixtures. In this study we generate acetate tolerant *P. putida* strains that overcome these challenges and examine their ability to produce isoprenol. We show that acetate tolerance and isoprenol production, though independent phenotypes can both be optimized in a given *P. putida* strain. Using proteomics and whole genome sequencing we examine the molecular basis of both phenotypes and show that tolerance to acetate can occur via alternate routes and result in different impacts on isoprenol production.

## Introduction

The 2023 United States Department of Energy “Billion Ton Report” confirmed that over 1.1 billion tons of plant-derived biomass is generated annually in the United States (1). This carbon stream, if harnessed, could provide an abundant source of renewable energy available to reduce greenhouse gas emissions from fossil fuels, decarbonizing the aviation, rail, and vehicle sectors (2) by upgrading simple carbon into valuable molecules through microbial bioconversion. While there are many routes to preparing monomeric carbon streams from plant biomass, the three major constituents extracted from the hemicellulosic and cellulosic fractions are glucose, xylose, and acetate (3–5). Glucose is already used in microbial bioconversion processes at scale to produce ethanol, now a gasoline blendstock prevalent in the automobile sector (6). However emerging molecules with advantaged properties could catalyze adoption of biobased processes in additional sectors, like aviation (7–10) or to generate plastics with improved recyclability (11). Utilizing as many types of carbon liberated from biomass has the potential to lower overall costs, and is imperative for an economically competitive process and commercial adoption (12, 13).

Of the two remaining major lignocellulosic-derived carbon sources, acetate (from xylan) can range from 0.5% to 17% (w/w) in concentration depending on the biomass source and pretreatment method utilized (14, 15) but is typically ignored as an undesirable byproduct of biomass pretreatment despite being a native substrate for many industrial hosts at lower concentrations. This could be attributed to acetate inhibiting microbial growth at higher levels and feed rates (12, 16). When protonated, acetic acid is lipophilic, enabling it to cross the cellular membrane, leading to cytoplasmic acidification and loss of membrane potential, contributing to its pleiotropic impacts on cell metabolism. Despite this toxicity, several reports have investigated the feasibility of using this organic acid primarily in *E. coli* as an alternative carbon source for the production of molecules such as itaconic acid (17), isobutanol (18) and succinate (19) and is comprehensively reviewed in (3).

Isoprenol (3-methyl-3-buten-1-ol) is a five carbon terpene molecule emerging as an advantaged biofuel and commodity chemical with demonstrated production across a range of microbial hosts using glucose as the carbon source, including *Pseudomonas putida* (20)*, Escherichia coli* (*21–23*), *Corynebacterium glutamicum* (24, 25) and budding yeast (26). Isoprenol can be used as a diesel-blendstock (27) or catalytically upgraded to 1,4-dimethylcyclooctane (DMCO), a sustainable aviation fuel (SAF) blendstock (13). In the case of *P. putida*, a microbe valuable for its ability to convert a range of carbon sources, inactivation of an isoprenol catabolism pathway was also needed to achieve bioproduction, in addition to a heterologous isoprenol production pathway, and isoprenol titers in *P. putida* KT2440 were improved upon deletion of PP_2675 (28), a cytochrome-C family PQQ regeneration cofactor needed for alcohol dehydrogenase activity (29).

Here we examine the initial *P. putida* ΔPP_2675 Δ14-PP_2676 strain (hereafter referred to as “PT”) response to growth on mixed glucose-acetate carbon streams. We took an approach that would enable us to identify mutants with improved acetate consumption that were also competent to produce isoprenol from mixed carbon stream conditions mimicking ratios found in hydrolysates (*e.g.,* ionic liquid pretreated hydrolysates) (30, 31). We were able to generate and optimize strains towards this goal and provide a deeper understanding of the underlying mechanisms using proteomics. The results of this study provide a foundation to co-utilize acetate from lignocellulosic biomass in an isoprenol conversion pipeline.

## Methods

### Strains and plasmids

All strains used in this study are listed in **Table 1**. A mutant strain of *Pseudomonas putida* KT2440 (ΔPP_2675 Δ14-PP_2676, referred to as “pre-tolerized” or “PT” throughout this manuscript) was used as the base strain for the tolerization described in this work. Whenever necessary, PT or *P. putida* KT2440 wild-type (WT) were used as controls as indicated.

**Table 1.** Strains used in this study. Whenever applicable, JBEI Inventory of Composable Elements (ICE) registry codes are provided in parentheses.

| Strain/Plasmid Name | Genotype | Notes | Source |
| --- | --- | --- | --- |
| WT | *P. putida* KT2440 | Wild type strain | ATCC 47054 (JBEI-18711) |
| PT | *P. putida* KT2440 ΔPP_2675 Δ14-PP_2676 | Pre-tolerization strain | (29) (JBEI-147164) |
| Σ1<sup>1</sup> | *P. putida* KT2440 ΔPP_2675 Δ14-PP_2676 Acetate<sup>+</sup> | Acetate/Isoprenol tolerized strain (Batch 2H) | This work (JBEI-253930) |
| Σ2 | *P. putida* KT2440 ΔPP_2675 Δ14-PP_2676 Acetate<sup>+</sup> | Acetate/Isoprenol tolerized strain (Batch 4P) | This work (JBEI-253931) |
| Σ3 | *P. putida* KT2440 ΔPP_2675 Δ14-PP_2676 Acetate<sup>+</sup> | Acetate/Isoprenol tolerized strain (Batch 5G) | This work (JBEI-253932) |
| Σ4 | *P. putida* KT2440 ΔPP_2675 Δ14-PP_2676 Acetate<sup>+</sup> | Acetate/Isoprenol tolerized strain (Batch 5H) | This work (JBEI-253933) |
| Σ5 | *P. putida* KT2440 ΔPP_2675 Δ14-PP_2676 Acetate<sup>+</sup> | Acetate/Isoprenol tolerized strain (Batch 5P) | This work (JBEI-253934) |
| pIY670 | pRK2-Kan-*araC*-P<sub>BAD</sub>-MvaS<sub>ef</sub>-MvaE<sub>ef</sub>-T<sub>rpoH</sub>-P<sub>tr</sub><br>c1-O-MK<sub>mm</sub>-PMD<sub>HKQ</sub>-AphA | Isoprenol pathway plasmid | Banerjee *et al*, 2024 (JBEI-233635) |
| pTE452 | pORTMAGE-Pa1 *recT* *mutL-E36K* *aaaC1* (gntR) | Recombineering plasmid | (32) Addgene # 138474, |
| pTE943 | *P<sub>LacM</sub>*-RBSopt-*aCpf1* *P<sub>J23119</sub>*-PP_0988-*gRNA*<br>LacI<sup>-</sup> BBR1 kanR | aCpf1 gene editing gRNA plasmid targeting PP_0988 (*gcvP-I*) | This work (JBEI-258301) |

<sup>1</sup> the Σ-class nomenclature was chosen to represent this class of tolerant mutants as $\Sigma$ is associated with summation, the idea of bringing together components of a set (akin to the utilization of many components from biomass), while the capitalized form of this greek letter also pays homage to the lowercase letter, as sigma factors ( $\sigma$ ) are a family of regulatory elements that regulate growth on mixed carbon sources.

|  |  |  |  |
| --- | --- | --- | --- |
| pTE944 | *P<sub>LacM</sub>*-*RBSopt-aCpf1 P<sub>J23119</sub>*-PP_2037-*gRNA*<br>LacI<sup>-</sup> BBR1 kanR | aCpf1 gene editing<br>gRNA plasmid<br>targeting PP_2037 | This work<br>(JBEI-258303) |
| pTE955 | *P<sub>LacM</sub>*-*RBSopt-aCpf1 P<sub>J23119</sub>*-PP_4264-*gRNA*<br>LacI<sup>-</sup> BBR1 kanR | aCpf1 gene editing<br>gRNA plasmid<br>targeting PP_4264<br>(*hemN*) | This work<br>(JBEI-258304) |
| Σ4 *ΔhemN*<br>GS-108 | *P. putida* KT2440 Σ4 ΔPP_2675<br>Δ14-PP_2676 Acetate<sup>+</sup> *ΔhemN*/ΔPP_4264 | Figure 5 | This work |
| Σ5 *ΔhemN*<br>GS-109 | *P. putida* KT2440 Σ5 ΔPP_2675<br>Δ14-PP_2676 Acetate<sup>+</sup> *ΔhemN*/ΔPP_4264 | Figure 5 | This work<br>(JBEI-261225) |
| Σ4 *ΔhemN*<br>GS-107 | *P. putida* KT2440 Σ4 ΔPP_2675<br>Δ14-PP_2676 Acetate<sup>+</sup> *ΔhemN*/ΔPP_4264 | Figure S5 | This work |
| Σ4 *ΔgcvP-I*<br>GS-089 | *P. putida* KT2440 Σ4 ΔPP_2675<br>Δ14-PP_2676 Acetate<sup>+</sup> *ΔgcvP-I*/ΔPP_0988 | Figure S5 | This work |
| Σ4 *ΔgcvP-I*<br>GS-090 | *P. putida* KT2440 Σ4 ΔPP_2675<br>Δ14-PP_2676 Acetate<sup>+</sup> *ΔgcvP-I*/ΔPP_0988 | Figure S5 | This work |
| Σ5 *ΔgcvP-I*<br>GS-093 | *P. putida* KT2440 Σ5 ΔPP_2675<br>Δ14-PP_2676 Acetate<sup>+</sup> *ΔgcvP-I*/ΔPP_0988 | Figure S5 | This work |
| Σ5 *ΔgcvP-I*<br>GS-094 | *P. putida* KT2440 Σ5 ΔPP_2675<br>Δ14-PP_2676 Acetate<sup>+</sup> *ΔgcvP-I*/ΔPP_0988 | Figure S5 | This work |
| Σ5 ΔPP_2037<br>GS-095 | *P. putida* KT2440 Σ4ΔPP_2675<br>Δ14-PP_2676 Acetate<sup>+</sup> ΔPP_2037 | Figure S5 | This work |
| Σ5 ΔPP_2037<br>GS-096 | *P. putida* KT2440 Σ4 ΔPP_2675<br>Δ14-PP_2676 Acetate<sup>+</sup> ΔPP_2037 | Figure S5 | This work |

Heterologous production of isoprenol was demonstrated using plasmid pIY670, which contains an optimized IPP-bypass pathway under regulation of an arabinose-induced promoter as well as the *neo* kanamycin selection marker (20). Deletion strains were generated following a previously described Cpf1-mediated ssDNA recombineering protocol (32–34). This recombineering plasmid was used in one tolerization batch (refer to the subsequent “tolerization” method section). Expanding the reported approach (35–37) to a genome-wide level, the frequency of spontaneous mutations was increased in each population by expressing the dominant negative *mutL-E36K* mutant, inhibiting the endogenous DNA mismatch repair system and enabling accrual of DNA mismatches primarily from replication (35–37).

For whole genome sequencing (WGS) of selected samples, genomic DNA was extracted from cell cultures using GenElute™ Bacterial Genomic DNA Kit (Sigma-Aldrich, St. Louis, MO) following the manufacturer’s protocol. WGS was conducted in 2×151bp paired-end reads runs in a Illumina NovaSeq X Plus sequencer by SeqCenter, LLC (Pittsburg, PA). Sequencing libraries were prepared using a tagmentation-based and PCR-based Illumina DNA Prep kit and custom IDT 10bp unique dual indices (UDI) with a target insert size of 280 bp. No additional DNA fragmentation or size selection steps were performed. Demultiplexing, quality control and adapter trimming was performed using Illumina’s bcl-convert (v4.2.4). Trimmed and filtered reads were mapped to the reference genome (NCBI SRA NC_002947.4) and in turn variant calls were identified using the software package Geneious (BioMatters LLC) with default parameters specified. Low quality variant calls adjacent to the ends of trimmed reads were filtered out by manual inspection after genome alignment.

### Culturing conditions

Routinely, cells were plated from cryostocks stored at -80 °C onto solid LB agar plates and incubated overnight at 30 °C. Single colonies were then picked and inoculated in 5 mL of liquid LB medium and incubated at 30 °C with agitation (200 rpm, orbital shaking) for 24 h without antibiotics, except as noted for the selection of plasmids. *P. putida* tolerization and the subsequent phenotypic characterization experiments were performed using minimal salt (M9) medium composed of 1x M9 salts (2 g/L (NH_4_)_2_SO_4_, 6.8 g/LNa_2_HPO_4_, 3 g/L KH_2_PO_4_, 0.5 g/L NaCl), 2 mM MgSO4, 0.1 mM CaCl_2_ and trace metal solution (500µL per 1L medium; Product No. 1001, Tekova Inc, Hollister, CA). Sodium acetate (Sigma-Aldrich, St. Louis, MO) was used in the composition of the M9 acetate medium. For all experiments, except when noted, acetate 50mM (∼4% w/v) or glucose 1 % (w/v) were used as the single sole carbon source in M9 medium. Acetate concentrations in media were calculated accounting for the contribution of the sodium fraction in the molecular weight of the sodium acetate salt. Isoprenol (3-methyl-3-buten-1-ol, 97% purity, 129402) was purchased from Sigma-Aldrich and added directly to the filtered M9 medium to specified concentrations as described in the text.

### Tolerization of PT to acetate and isoprenol

For generating acetate and isoprenol tolerant isolates in the *P. putida* PT strain background, a single colony was first picked from a fresh agar plate and cultured in LB medium in the conditions described above. The following day, the saturated culture was back diluted (1:100) in 25 mL of M9 acetate in 250mL baffled shake flasks for an initial acetate acclimation step. Cells were allowed to grow in M9 acetate in the conditions described above for up to 24 h to overcome lag phase, then adapted cells were harvested by centrifugation at 4,000 x g for 10 minutes at room temperature and resuspended in 500 μL - 1000 μL of fresh M9 acetate. The resulting concentrated cell suspension was diluted to a target OD_600_ of 0.01 in different flasks containing 25 mL of M9 acetate supplemented with isoprenol (7 g/L, 8 g/L and 9 g/L). 2mL aliquots of these new cultures were then distributed in 24-well deep-well plates, which were sealed and incubated at 30 °C with agitation. Plates were left in the incubator for up to five days and regularly visually inspected for turbidity in the medium.

As soon as cultures became turbid (usually two to three days after inoculation), an aliquot was spread on LB agar for retrieving isolated colonies. A single colony from each new plate was then again grown under the tolerization conditions, starting from the initial LB cultivation, and stressed in M9 acetate medium supplemented with isoprenol (8 g/L and 9 g/L) for validation of the tolerance phenotype. Cultures that grew under the validation conditions were then once again back diluted in M9 acetate medium supplemented with isoprenol (10 g/L and 11 g/L) as an additional filter against false positives (clones that resumed growth only after isoprenol evaporation). Single colonies derived from cultures that grew robustly under the screening conditions were then retrieved and stocked with glycerol (25% w/v final) for phenotypic characterization.

We considered each individual starter LB culture as an experimental batch of tolerization. For the batch with cells expressing the recombineering plasmid, an overnight culture of cells freshly transformed with pORTMAGE Pa-1 (32), was diluted 1:100 in 5 mL of fresh LB medium and allowed to outgrow for one hour. After this, recombinase expression was induced with 1 mM 3-methyl-benzoate (Sigma Aldrich T36609; m-toluic acid; “3-MB”), and cultures were allowed to resume growth overnight before being used in the tolerization experiment (as described earlier). In these cultures, gentamicin (30 μg/mL) was added for plasmid maintenance, but the antibiotic was not used in the follow up validation experiments, and all isolates retrieved from this batch were verified to be gentamicin sensitive during the subsequent phenotypic characterization stages.

### Measurement of growth kinetics

Growth profiles of cells cultivated in different conditions were assessed in microtiter well plate format. In these experiments, given the different performances of PT, WT, and the tolerant isolates when adapting from rich medium to M9 acetate medium, the saturated LB cultures were first back diluted (1:100) in 5 mL of M9 glucose medium and grown overnight. The following day, saturated cultures were diluted to a final OD_600_ of 1.0/mL in 500 μL of M9 medium without carbon sources. These cell suspensions were finally diluted (1:100) in 300 μL of fresh medium in 48-well microplates. The plates were sealed with semi-permeable films (Breathe Easy Film, USA Scientific, Ocala, FL) and incubated for 48 hours in a microplate reader (Molecular Devices m2E Plate Reader, San Jose, CA) with continuous agitation and temperature controlled at 30 °C. Readings were taken every 15 minutes. Experiments were performed with at least three biological replicates and on different days from different starting cultures, and at least two technical replicates (repeated wells) per strain and condition. Microplate reader data was analyzed using an in-house workflow in R (mipreadr v0.1, available at github.com/guazzaroniLab/mipreadr).

### Proteomics analysis

Fresh colonies were inoculated in 5mL of LB and cultured under the standard conditions described above. After the M9 glucose adaption, cells were transferred to 50 mL conical tubes and harvested by centrifugation for 5 minutes at 4000 rpm. The cell pellet was washed twice with 1000 µL of M9 minimal medium without carbon sources and resuspended in 800 μL of M9 medium without carbon sources. Each of these initial cell suspensions was used to inoculate three cultures of either M9 Acetate, M9 Glucose or M9 Glucose-Acetate at an initial OD_600_ 0.15 in 5 mL cultures. After inoculation, cells were incubated at 30 °C with agitation (200 rpm) and microbial growth was periodically tracked by the measurement of turbidity. Once cultures reached exponential growth (OD_600_ of 0.6 - 0.8), the cultures were transferred to new 50 mL conical tubes and the cells harvested by centrifugation at 4000 *xg* for 5 minutes. Finally, the cells were resuspended in 100 μL of M9 medium without carbon sources and harvested in PCR strips that were stored at –80 °C until further processing. Protein was extracted from cell pellets and tryptic peptides were prepared by following an established proteomic sample preparation protocol (38). Peptide samples were loaded onto an Ascentis® ES-C18 Column (Sigma–Aldrich, USA) and were eluted from the column by with a 10 minute gradient from 98% solvent A (0.1 % FA in H_2_O) and 2% solvent B (0.1% FA in ACN) to 65% solvent A and 35% solvent B. Eluting peptides were introduced to the mass spectrometer operating in positive-ion mode and were measured in data-independent acquisition (DIA) mode with a duty cycle of 3 survey scans from m/z 380 to m/z 985 and 45 MS2 scans with precursor isolation width of 13.5 m/z to cover the mass range. DIA raw data files were analyzed by an integrated software suite DIA-NN (39). The databases used in the DIA-NN search (library-free mode) was *P. putida* KT2440 latest Uniprot proteome FASTA sequences (generated in March 2024) and common proteomic contaminants. The retention time extraction window was determined individually for all MS runs analyzed via the automated optimization procedure implemented in DIA-NN. Protein inference was enabled, and the quantification strategy was set to Robust LC = High Accuracy. The output main DIA-NN reports were filtered with a global FDR = 0.01 on both the precursor level and protein group level. The Top3 method, which is the average MS signal response of the three most intense tryptic peptides of each identified protein, was used to calculate the total intensity of proteins in the samples (40, 41). The cluster analysis (Figure 5C) was made with a subset of “high confidence” proteins, considering an arbitrary threshold of at least ≥300,000 protein counts by MS after top3 peptide filtering for at least one of the samples, prior to relative abundance normalization. DIA-NN is freely available for download from https://github.com/vdemichev/DiaNN.

### Substrate consumption measurements

Fresh colonies were inoculated in 5mL of LB and cultured under the standard conditions described above. The following day, cells were back diluted 1:100 in M9 glucose 2% (w/v) and allowed to grow for an additional period of 24 h. Cells from these saturated cultures were then harvested by centrifugation for 5 minutes at 4000 *xg* and washed twice using 1 mL of M9 minimal medium without carbon sources. Washed cells were resuspended in 800 uL of M9 medium without carbon sources and inoculated to an initial OD_600_ of 0.2 into three separate tubes of either M9 glucose 2% (w/v), M9 acetate 50mM (4% w/v), or M9 glucose 2% (w/v) with acetate 0.65% (w/v) in 5mL cultures mimicking mixed carbon concentrations from complex sorghum hydrolysates (42). Cells were grown at 30 °C with agitation in an orbital shaker at 200 rpm. Aliquots of the culture media were taken immediately after inoculation (0h), and at points 6h, 12h, 18h, 24h and 48h post inoculation. Cell growth was measured via turbidity (OD_600_) and 200 μL aliquots of each timepoint were stored at –20 °C before the determination of acetate and glucose concentrations in the supernatant via HPLC. Residual glucose and acetate were analyzed using an Agilent 1260 HPLC (Agilent Technologies, USA) equipped with a refractive index detector (RID) and an Aminex 87H column in an oven maintained at 60 °C. 4 mM sulfuric acid was used as the mobile phase at 0.6 mL min^-1^. Substrate concentrations were quantified based on standard calibration curves generated using authentic standards for glucose and acetate (Sigma Aldrich, USA).

### Isoprenol production runs

Fresh colonies of strains of the indicated genotype transformed with the isoprenol production plasmid pIY670 (20) were inoculated in 5mL of LB medium in the test tube format containing 50 μg/mL kanamycin and cultured at 30 °C and 200 rpm and handled as described in (20) with modifications outlined below. Briefly, saturated overnight cultures were back diluted (1:100) in M9 glucose 2% (w/v) and cultured for an additional day for minimal media adaptation. For isoprenol production runs, cells were harvested by centrifugation for 5 minutes at 4,000 *xg* and washed twice using 1 mL of M9 minimal medium without carbon sources. The washed cells were resuspended in 800 µL of M9 minimal medium without carbon sources and back diluted to an initial OD_600_ of 0.2 in 5mL of fresh M9 medium with glucose 2% (w/v) or glucose 2% (w/v) with acetate 0.65% (w/v) in triplicates. Arabinose was added at a final concentration of 0.2% (w/v) in the medium to induce the production pathway, and kanamycin 50 μg/mL was used for plasmid maintenance. Cells were grown at 30 °C with agitation. Isoprenol production was monitored at the 24h, 48h and 72h post inoculation timepoints to measure cell growth via turbidity (OD_600_) and isoprenol titer by harvesting 200 µL cell culture aliquots

To determine the concentration of isoprenol in the samples, samples were harvested and prepared for an ethyl acetate extraction method as described previously (26, 43) for quantification with a gas chromatography-flame ionization detector (GC-FID, Agilent Technologies) fitted with an Agilent DB-WAX capillary column (15 m x 320 μm x 0.25 μm). The GC method used was executed exactly as previously described and data was acquired and analyzed using OpenLab (Agilent Technologies). Authentic isoprenol (TCI America, Catalog No. M0726) was used to generate a calibration curve for quantification.

## Results

### The Initial Isoprenol Catabolism-Deficient *P. putida* Strain Shows Arrested Growth in Media Containing Acetate

Glucose and acetate are native substrates that support biomass formation in *P. putida* KT2440 when supplied separately and as a mixed substrate. However, we found that WT *P. putida* shows an impaired growth profile compared to either single carbon stream condition (**Figure 1A**). As our laboratory had already developed strains deficient for isoprenol catabolism (29), we characterized the behavior of ΔPP_2675 Δ14-PP_2676 strain (referred to as “PT”) for growth on the same carbon streams to assess the feasibility of biofuel production in alternative growth regimens. Our results show that, in all cases, the PT strain has a slower doubling time than the WT control; while growth on glucose is similar to the WT, on the sole acetate medium we observe an extended lag phase before a detectable turbidity increase roughly delayed by 12 hours compared to WT. Consistent with these observations, the substrate consumption profiles of WT and PT for glucose as a single carbon source are similar (**Figure 1B**), and the increased lag time for the PT strain when grown on acetate correlates with the delayed timing in observed decrease in acetate concentration (**Figure 1C**). Adjusted linear models for the substrate consumption rate indicate that within the first 24 hours of incubation, WT consumes glucose at a rate of 0.846 h^-1^ (R^2^ = 0.975), whereas for PT this rate is 0.824 h^-1^ (R^2^ = 0.977). This suggests that, despite PQQ being a cofactor needed for the assimilation of glucose via glucose dehydrogenase (44), PP_2675 is not required for glucose catabolism in the M9 glucose condition. In contrast, the growth and consumption profiles of the PT strain are markedly different in the dual carbon source experiment. While both glucose and acetate are co-utilized simultaneously at similar rates in WT in the sole glucose or sole acetate conditions, the overall consumption rate for both glucose or acetate decreases from 0.824 h^-1^ (R^2^ = 0.977) to 0.262 h^-1^ (R^2^ = 0.975) for glucose and 0.20 h^-1^ (R^2^ = 0.870) to 0.058 h^-1^ (R^2^ = 0.778) for acetate in PT (compare **Figure 1B, 1C** to **1D**). The requirement for PQQ cofactor regeneration is apparent under the mixed glucose-acetate condition, suggesting an important role for PP_2675 in this substrate and its possible relevance to isoprenol production with acetate-containing carbon streams.

**Figure 1.**
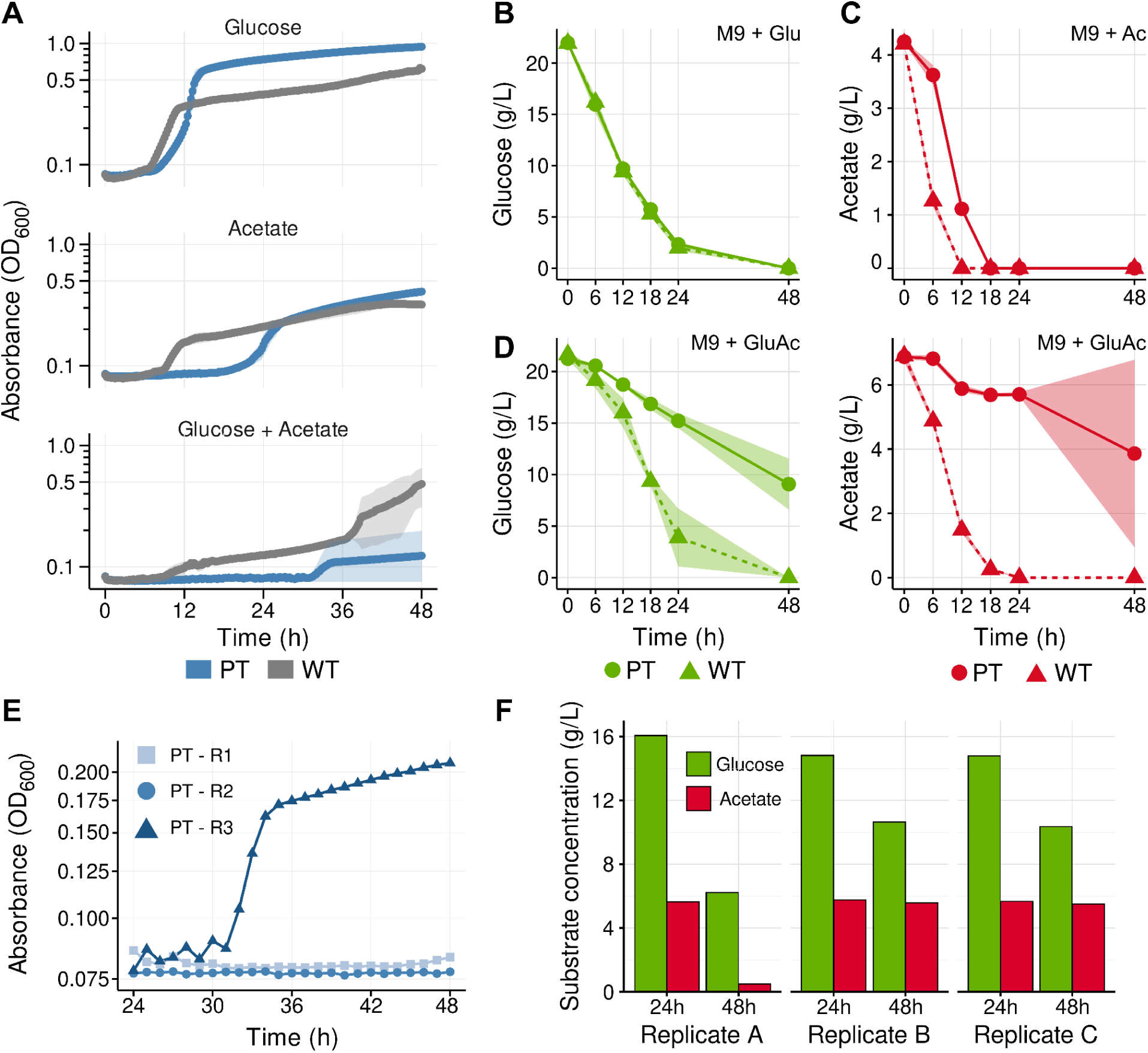
Phenotypic characterization of *P. putida* WT and PT strains grown in minimal media containing glucose, acetate or a mixture. A) kinetic growth curves of *P. putida* KT2440 Wild Type (WT) and KT2440 ΔPP_2675 Δ14-PP_2676 (“PT”) in minimal M9 media supplemented with the carbon sources glucose 1% (w/v) (top), acetate 4% (w/v) (middle) or a mixture (model hydrolysate) containing glucose 2% (w/v) and acetate 0.65% (w/v) (bottom) in microtiter dish format. **B)** Measurement of residual glucose concentrations in the supernatant of cultures grown in test tubes with glucose-only medium. **C)** Measurement of residual acetate concentrations in the supernatant of cultures grown in test tubes with acetate-only medium. **D)** Measurement of residual glucose (left) and acetate (right) in the supernatant of cultures grown in test tubes with model hydrolysate medium. **E)** Growth profile of the individual biological replicates of PT (R1, R2 and R3) displayed in **A** after 24 hours of growth in M9 acetate + glucose. **F)** Glucose and acetate consumption profile of the individual biological replicates of PT (Replicate A, Replicate B and Replicate C) displayed in **D** at timepoints 24 and 48 hours after the beginning of the experiment. The shaded area surrounding the line represents the standard deviation of measurements between 3 biological replicates for all carbon in panels **A**, **B**, **C**, and **D**.

The averaged values for growth indicated high variability between PT replicates in the mixed carbon source experiments. Re-examining the biological replicates separately indicated that only one out of three replicates was able to grow (**Figure 1E**). A similar result was observed in the consumption experiment, where, despite residual glucose concentrations noticeably decreasing in all three replicates, only a subset of the biological replicates were able to resume growth in this mixed carbon source media (**Figure 1F**). In contrast, all replicates in the WT strain showed similar growth with some variability after 36 hours incubation. In both cultivation formats for PT, the single replicate that grew exhibited an extended lag phase (>30 hours), longer than when grown in acetate as a sole carbon source. It is possible that basal levels of glucose consumption in the dual carbon source condition allows PT cells to remain viable during their long lag phase, despite robust growth only being possible once acetate is entirely depleted from the medium. Accordingly, the final OD_600_ of PT culture that showed growth in the mixed carbon medium is much lower than the WT control and are similar to the levels reached in acetate-only medium by all strains (**Supplemental Figure S1**).

### Tolerization Allowed the Recovery of Strains with Improved Fitness in Acetate as a Carbon Source

Given the growth phenotype exhibited by PT in the mixed glucose-acetate carbon source minimal medium (**Figure 1A**), we looked to identify bypass mutants that showed improved growth in acetate-containing media by cultivating PT cells in media containing both acetate and growth inhibitory concentrations of isoprenol. By also including growth-inhibitory concentrations of isoprenol in the growth medium, we reasoned that this stringent growth threshold would select for mutants with improved fitness in acetate as a carbon source that were also tolerant to the desired end product (**Materials and Methods**). Isoprenol concentrations above 4 g/L are inhibitory for cell growth of WT *P. putida* in deep well plates, and over the course of three months, we screened PT clones isolated from M9 acetate supplemented with 7-9 g/L isoprenol in five independent experimental batches. Our workflow is in this case similar to a mutant recovery approach, as in each experimental batch we were able to recover isolates with improved phenotypes compared to PT (**Figure 2**). We refer to the validated set of acetate/isoprenol tolerant clones as the Σ-class strains (alluding to the use of the Greek letter “Σ” now commonly used as a term for summation) and numbered according to the chronological order in which they were added to our internal accession database. **Supplemental Table 1** summarizes the screening conditions in which each of the Σ-class strains were isolated.

**Figure 2.**
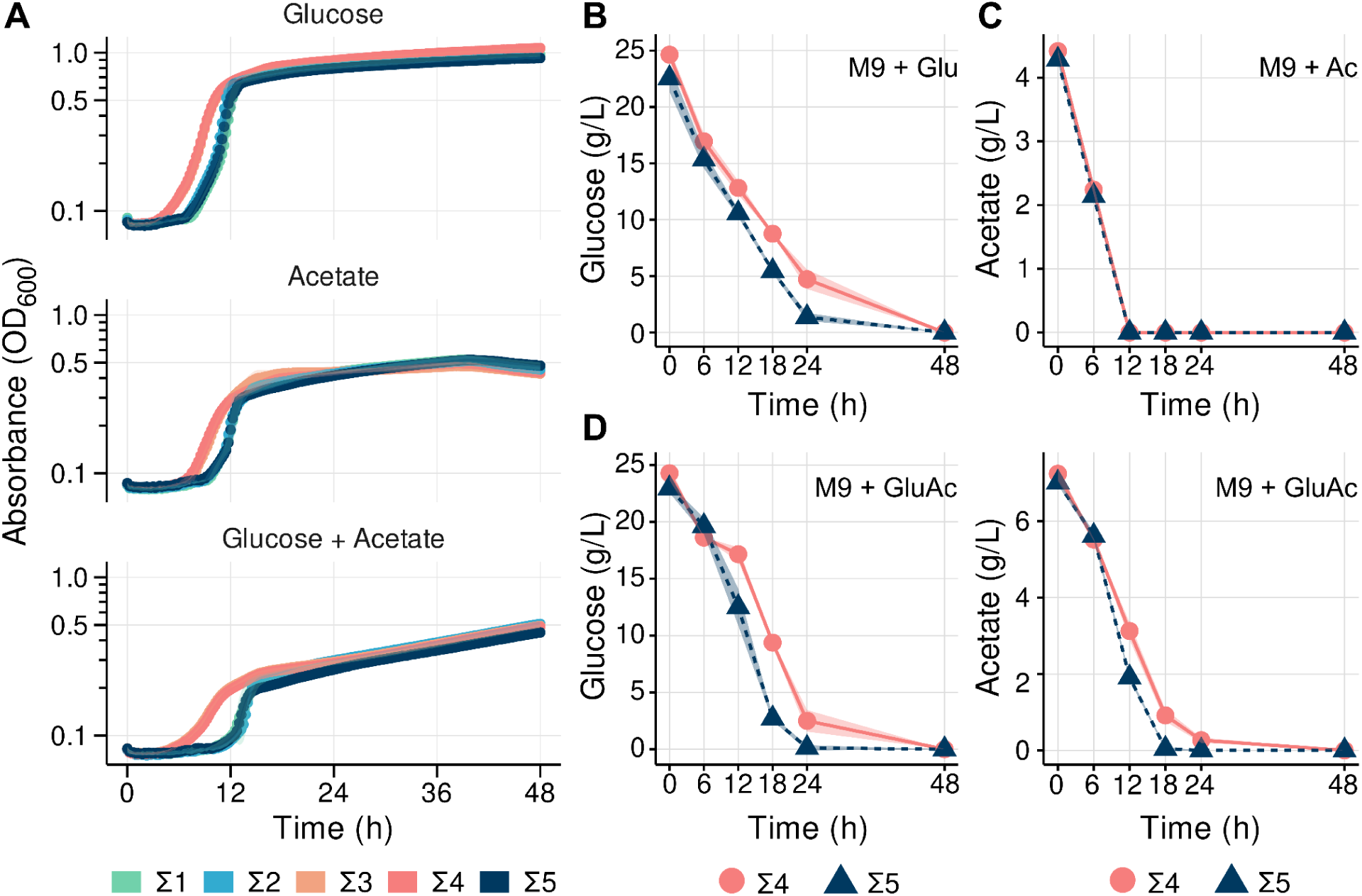
Phenotypic characterization of tolerized Σ-class strains in minimal media containing glucose, acetate or a mixture. **A)** kinetic growth curves of tolerized strains (Σ1, Σ2, Σ3, Σ4, Σ5) in minimal M9 media supplemented with the carbon sources glucose 1% (w/v) (top), acetate 4% (w/v) (middle) or a mixture (model hydrolysate) containing glucose 2% (w/v) and acetate 0.65% (w/v) (bottom) in the microliter dish format. **B)** Measurement of residual glucose concentrations in the supernatant of Σ4 and Σ5 cultures grown in test tubes with glucose-only medium. **C)** Measurement of residual acetate concentrations in the supernatant of Σ4 and Σ5 cultures grown in test tubes with acetate-only medium. **D)** Measurement of residual glucose (left) and acetate (right) in the supernatant of Σ4 and Σ5 cultures grown in test tubes with model hydrolysate medium. The shaded area surrounding the line represents the standard deviation of measurements between 3 biological replicates.

All Σ-class strains possess a shorter lag phase (reduced from ∼48 hours to 6 hours) when compared to the pre-tolerized strain in M9 acetate medium, as well as a faster maximum growth rate during the exponential phase (**Figure 2A**, refer to growth of strain PT in **Figure 1**). We also highlight the reproducibility of strain growth where all biological replicates show rapid entry into the exponential phase at comparable timing to WT, compared to the highly variable growth in the PT strain (**Figure 1**). Despite the improvements in growth in acetate and glucose-acetate when compared to the parental strain, Σ-class strains perform slightly worse in acetate medium supplemented with as little as 2 g/L isoprenol in comparison to acetate-only medium, although growth rates and lag phase durations remain superior than what is observed for PT (**Supplemental Figure S2**).

Among the five Σ-class strains, we observed two phenotypic groups: the Σ3 and Σ4 strains show a more rapid adaptation to media containing either acetate, glucose or a mixture, and a second group comprised of the Σ1, Σ2 and Σ5 strains, which have a more noticeable lag phase in medium containing both acetate and glucose. To better understand the two growth dynamics, we performed a substrate consumption time course for representative strains Σ4 and Σ5 in minimal media containing different carbon sources (**Figure 2 B-D**). In the mixed carbon source media, during the first 24 hours, consumption of glucose by Σ4 was slower (0.879 h^-1^, R^2^ = 0.949) than for Σ5 (1.04 h^-1^, R2 = 0.956), but acetate consumption occurred at similar rates (0.30 h^-1^ and 0.32 h^-1^ respectively). Despite these slight differences in substrate consumption kinetics, both Σ4 and Σ5 strains showed complete consumption of both glucose and acetate, suggesting the limitation imposed by the ΔPP_2675 mutation in PT strain had been remedied. Their robust growth rates and complete substrate utilization provided ideal characteristics for a biofuel bioconversion host, satisfying the initial goal of this study.

### Tolerization endowed cells with different isoprenol production capacities in minimal media

The next objective was to determine if the Σ-class strains were advantaged over the PT strain in producing isoprenol under different carbon stream compositions. We transformed a plasmid-based system harboring the isoprenol production pathway (**Materials and Methods**) into all Σ-class strains along with the parent PT strain and prepared isoprenol production runs in M9 minimal salt media with either glucose and acetate as sole carbon sources, as well as a glucose-acetate mixed carbon source. In M9 glucose medium, the base PT strain produced 283 ± 26 mg/L isoprenol by the 48 hour timepoint, but still showed substantial (35%) isoprenol loss by the 72 hour time point, suggesting this strain still retained some ability to degrade isoprenol (**Figure 3A**). Tolerized strains Σ3 and Σ4 showed slightly lower isoprenol titers in comparison to PT, the maximum isoprenol titers for these strains being 207 ± 22 mg/L and 205 ± 20 mg/L, respectively, while Σ1, Σ2, and Σ5 isolates showed increased isoprenol titers reaching up to 389 mg/L by the 48 hour time point. We note that Σ5 stain exhibits reduced isoprenol loss in the third day of production, with only a 2.73% loss in comparison to the previous 24 hours. This glucose to isoprenol titer improvement was not the objective of our tolerization effort, but underscores how a more complete characterization of strains can reveal unintended phenotypic changes.

**Figure 3.**
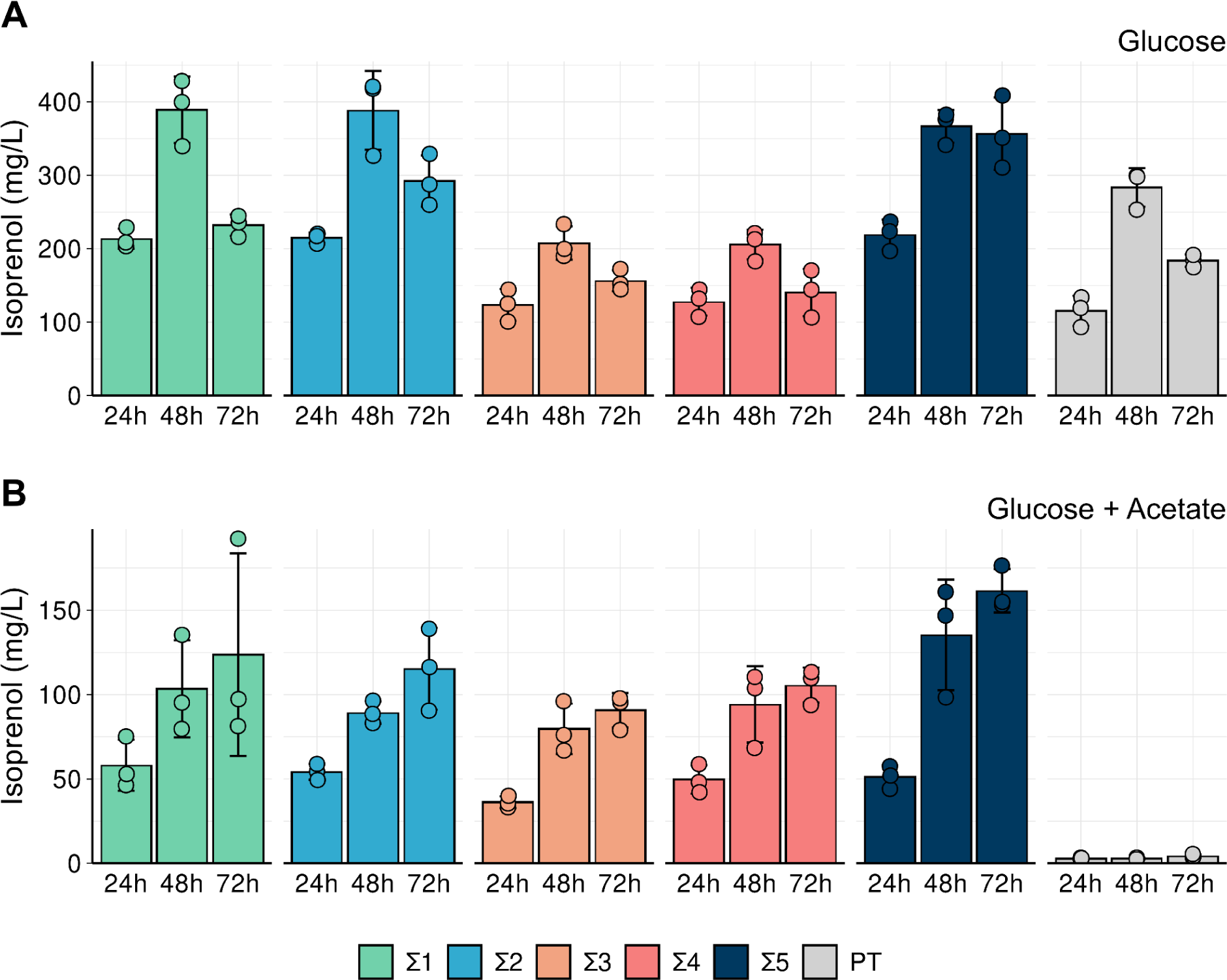
Plasmid-based production of isoprenol in PT and Σ-class strains. **A)** on glucose 2% and **B) on** glucose 2% (w/v) and acetate 0.65% (w/v). Samples were harvested at the indicated time points post isoprenol pathway induction with 0.2% arabinose. Datapoints shown represent 3 independent biological replicates, and the error bars indicate standard deviation from the mean.

When strains were cultivated in the glucose-acetate mixed carbon condition, we noted that the overall titers were lower than with the glucose alone condition, even though the overall supplied utilizable carbon was higher as the acetate concentration was not reduced in this cultivation medium (**Supplemental Table S2**, **Figure 3B**). In this production regime, the baseline PT strain failed to grow, and therefore did not produce any detectable isoprenol. In contrast, all Σ-class strains were able to produce isoprenol with titers at the 48 hour time point being comparable to production at the 72 hour time point for all strains. Σ5 reached the highest final isoprenol levels (161 ± 12 mg/L) at 72 hours. Acetate alone did not result in any detectable isoprenol production in Σ-class strains (**Supplemental Figure S3**), suggesting that acetate was used to support biomass formation and isoprenol formation was driven primarily from glucose as the carbon source. These results indicated that our initial selection regime provided a rapid strategy to identify PT mutant strains which could grow on the glucose-acetate mixed carbon source and were competent to produce isoprenol. Overall, the variable growth phenotypes of the Σ-class strains correlated with different production phenotypes, but, contrary to our expectations, titers for the tolerized strains with shorter lag phases were generally lower than for those with longer lag phases.

### Proteomics reveals different paths for acetate and glucose assimilation in tolerized and naive *P. putida* strains

To better assess how tolerization enabled different growth and production phenotypes across our strains, we investigated the proteomes of cells grown in acetate, glucose and glucose-acetate (**Figure 4**, **Figure 5**). Mean protein counts and relative protein abundances for each sample, as well as and other summary statistics for the proteomics dataset, are available in **Supplemental File 1**.

**Figure 4.**
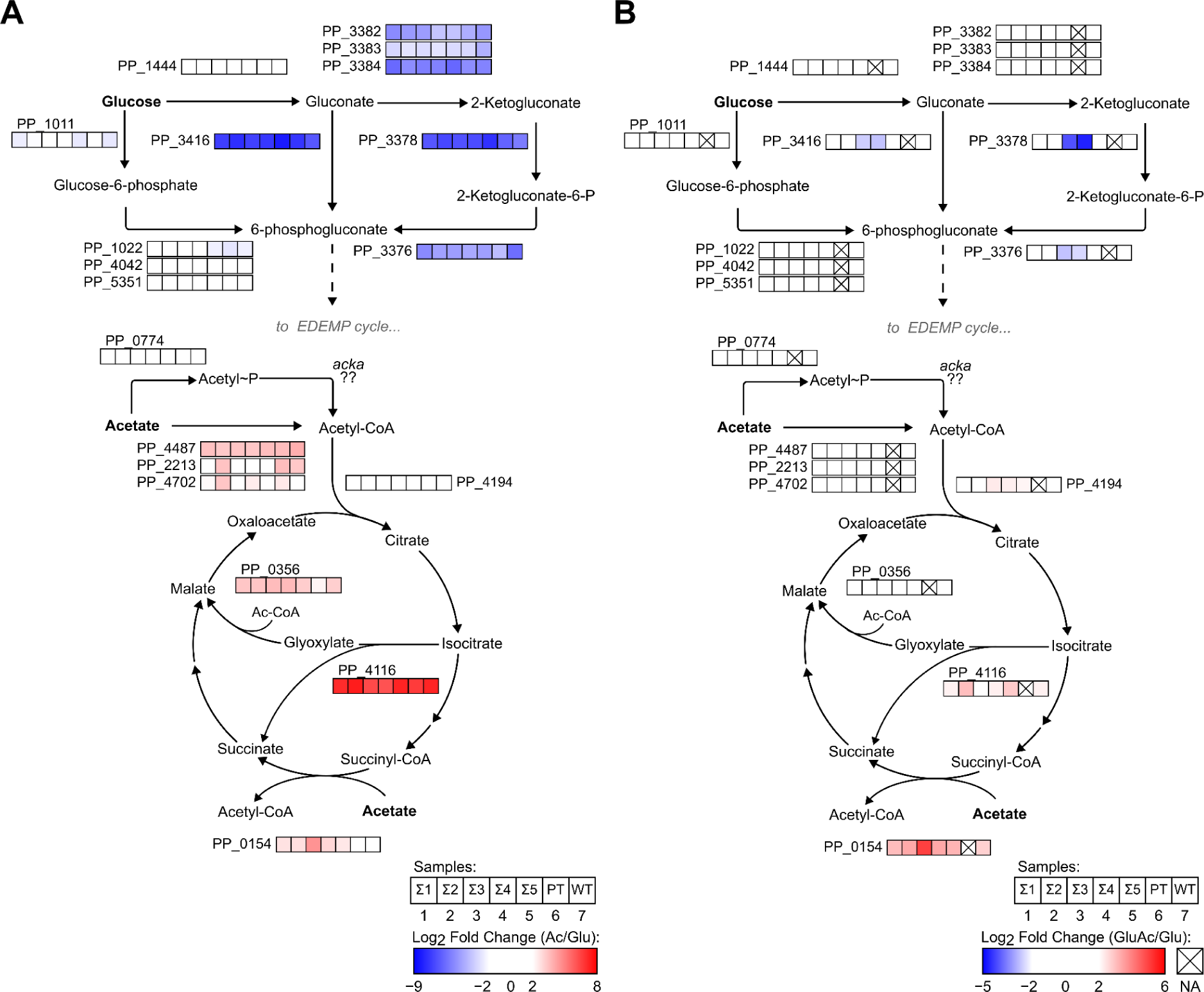
Differences in fold-change levels of proteins involved in glucose and acetate assimilation routes. Panels show differences in the expression levels (log2 fold change) of proteins differentially expressed in **A)** acetate as a sole carbon source and **B)** glucose and acetate as a dual carbon source compared against itself and cultivated in either medium. For both panels, protein levels in the glucose as a sole carbon source medium was used as the reference condition.

First, a differential protein expression analysis for cells grown in acetate in comparison to glucose (**Figure 4A**) suggested acetate utilization in PT was rate-limited in key steps in comparison to the Σ-class strains, including acetate assimilation and activity of the glyoxylate shunt. In acetate medium, the protein encoded by PP_0154 (SpcC) increased its expression at least twofold higher in all Σ-class strains, but not in PT and WT. This protein is a putative succinyl-CoA:acetate CoA-transferase by homology and motif analysis that allows acetate to enter the TCA cycle via succinate (45, 46). Energetically, the difference in how acetate is incorporated into the central carbon metabolism might lead to important differences in the availability of intracellular energetic currency, which might contribute to the faster growing phenotypes of Σ-class strains. Accordingly, it is possible that PT and WT underperform in acetate due to the incorporation into central metabolism by direct production of acetyl-CoA, a reaction that requires ATP (46).

This analysis also revealed that the proteins responsible for the incorporation of glucose into the central carbon metabolism via transformation of gluconate, e.g. gluconate kinase (PP_3416) and 2-ketogluconate kinase (PP_3378), are highly repressed (log2 fold change < 8) in acetate in comparison to glucose, but glucokinase, encoded by the gene PP_1011, remains at relatively similar absolute protein quantification expression levels in both conditions (the median value of protein counts for PP_1011 in glucose is 1.1 x 10^6^ and in acetate 2.7 x 10^5^. For PP_3378 the median values are, respectively 1.8 x 10^6^ and 8 x 10^3^). The fact that the presence of acetate in the medium does not promote a strong down-regulation in this pathway might explain why, despite failing in consuming acetate in the mixed carbon source medium, glucose levels in PT still decrease over time, allowing cells remain viable for long periods of time (**Figure 1**).

The difference in activation of the gluconate pathway is consistent with the phenotypic profiles observed for strains Σ4 and Σ5 regarding substrate consumption and isoprenol production (**Figure 2** and **Figure 3)**. In the glucose-acetate condition in comparison to the glucose only medium feed (**Figure 4B**), we observed that this pathway remains strongly repressed in Σ3 and Σ4 but not in Σ1, Σ2, Σ5 and WT, suggesting that the ability to use alternative routes for glucose consumption in the presence of acetate is an important factor for biofuel bioproduction in model hydrolysate medium. In addition to the differences in the uptake of substrates, a noticeable up-regulation in isocitrate lyase (PP_4116), the first step of the glyoxylate shunt, for strains Σ2 and Σ5 (log_2_ fold change of 3.01 and 2.82, respectively), can also have provided additional fitness benefits that reflect in better resource utilization by these strains. As PT cannot grow reliably in mixed carbon source medium without toleration, we could not characterize this strain in the glucose-acetate condition, but together, these results shed light on how phenotypic differences in acetate growth rates and isoprenol production can be realized through different proteomic responses across mutants in an acetate-glucose mixed feed regime impacts metabolic processes and has downstream impacts on using lignocellulosic substrates for bioconversion processes.

### Transcription Factors and Other Non-metabolic genes play important roles in the tolerance phenotype of Σ-Class strains

Despite the differences found regarding central metabolism in acetate medium, 78% of the proteins found to be differentially expressed by any Σ strain in comparison to PT that met the threshold for statistical significance did not belong to an annotated KEGG metabolic pathway, which suggests that other kinds of molecular effectors might be promoting tolerance phenotypes. To capture the overall complexity of this dataset we performed a principal component analysis (PCA) of all quantified proteins in the acetate growth condition, which indicated clustering of strains along the axis of larger variance that coincides with lag phase duration order in acetate-containing media observed in our phenotypic characterization experiments (**Figure 5A**). In this axis, slower growing strains (PT and WT) are grouped in one end of the plot, whereas faster growing strains (Σ3 and Σ4) were grouped in the opposite direction. Enzymes like the previously mentioned putative succinyl-CoA:acetate CoA-transferase PP_0154 (SpcC), succinate dehydrogenase (PP_4192), acetolactate synthase (PP_3365), and glutamate dehydrogenase (PP_0675) correlate with more positive values, while stress-tolerance proteins like CspD (PP_4010), AhpC (PP_2439) and the serine proteases ClpA (PP_4008) and ClpX (PP_2301) point to the opposite direction. CsrA (PP_3832) a post-transcriptional regulator known for acting on many cellular processes, including oxidative stress survival in bacteria (47) was also associated with longer lag phases in the acetate medium. This global analysis supported the notion that many diverse regulatory changes were likely driving improved growth on acetate in the Σ-class strains.

**Figure 5.**
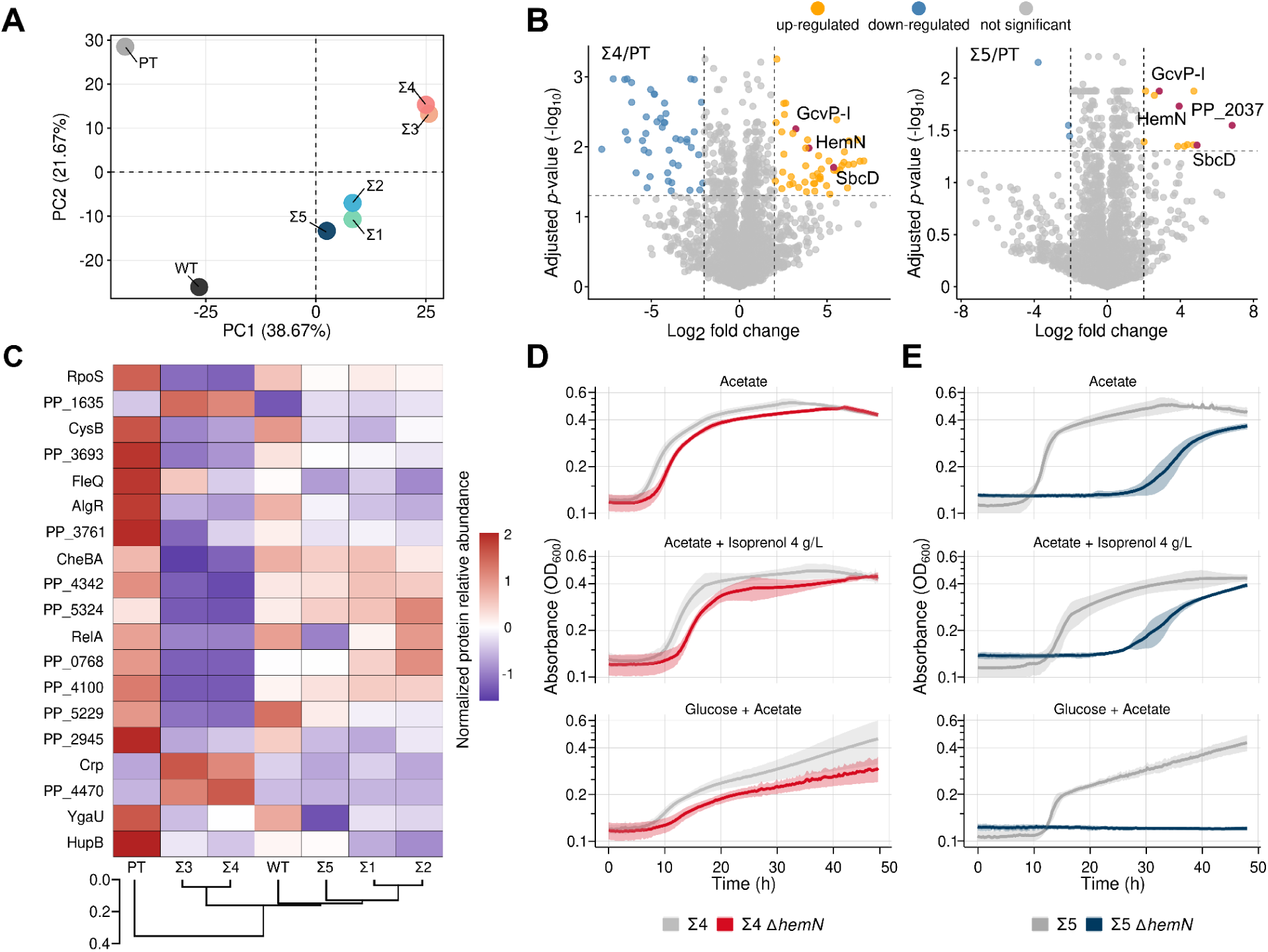
Global regulatory effectors contribute differently to acetate utilization and isoprenol production in different strain backgrounds. **A)** Principal component analysis on the proteomic profiles of strains grown in acetate as a sole carbon source. Points represent each of Σ-class, PT and WT strains plotted along the two directions of largest variance (PC1 and PC2). Relative protein abundances of proteins were used for dimensionality reduction. **B)** Volcano plots for protein levels of Σ4 (left) or Σ5 (right) in comparison to PT in acetate. Comparisons with BH-adjusted *p*-values (cutoff set to ≤ 0.05) and absolute log_2_ fold change >2 or <-2 were considered both statistically significant and differentially expressed. **C)** Heatmap highlighting differences in expression levels for different regulators of strains grown in acetate as the sole carbon source. Heatmap rows and columns are ordered based on a hierarchical clustering of the percent relative abundance values of each protein in its sample, using Euclidean distances as the dissimilarity metric. Z-score scaling was used to ensure values were on range for visualization, but not for clustering The distance between strain clusters is represented as a dendrogram plotted on the bottom of the panel. **D, E)** Phenotypic characterization of *hemN*/PP_4264 deletion in Σ4 (**D**) and Σ5 (**E**) in the conditions acetate 4% (w/v) glucose 2% (w/v) and acetate 0.65% (w/v), and acetate 4% (w/v) supplemented with isoprenol 4 g/L. The color-shaded area surrounding the solid line indicates the standard deviation from the mean from at least three independent biological replicates.

Next, we performed pairwise comparisons of protein expression levels between the Σ-class strains to the PT strain to identify the specific protein effectors implicated by a differential response to acetate. We cross-referenced proteins to their functional genomic characteristics in the RB-TnSeq fitness database (48) and known functions from the literature to identify potential activities related to acetate catabolism or strain fitness. For instance, the acetate permease ActP-I (PP_1743), was found to be downregulated in all Σ-class strains but Σ5. Additionally, strains Σ3 and Σ4, which belong to their separate PCA cluster also exhibited downregulation of a surface adhesion protein (PP_0168) and upregulation of an uncharacterized transcriptional regulator (PP_0673) as well as a strong downregulation (log_2_ Fold Change < 5) in RelA (PP_1656), which are phenotypes consistent with mild fitness improvements in the medium with acetate as a sole carbon source according to RB-TnSeq data. We show the distribution of differential protein expression of the representative Σ4 and Σ5 strains compared to the PT strain grown in the acetate medium using volcano plots (**Figure 5B**). For all Σ-class strains, the oxidoreductase-family glycine decarboxylase encoded by PP_0988 (GcvP-I), the subunit D of the nuclease SbcCD encoded by PP_2025, the coproporphyrinogen-III oxidase encoded by PP_4264 (HemN) and an UPF0313 protein encoded by PP_4872 (YgiQ) are all significantly overexpressed in comparison to PT in the acetate medium. In this condition, PP_2037, a putative aldolase, is also up-regulated in Σ1, Σ2 and Σ5, the cluster of Σ-class strains with better isoprenol production. While little information is available for this protein, PP_2037 has been reported to be differentially expressed in a *P. putida* strain mutant for FinR, an oxidative stress-sensing transcriptional regulator in proteobacteria (49).

The differential proteomics analysis identified key protein effectors that likely mediate the increased acetate tolerance in Σ-class strains. This analysis was complemented by whole genome resequencing of parental PT strain and four representative Σ-class strains in an attempt to identify causal mutations driving the global cellular rewiring. However, little overlap was observed in mutated ORFs or intergenic regions identified by the DNA sequencing and the differential expression targets identified off of the proteomic data (**Supplemental Figure S4, Supplemental File 2**). While alterations in expression levels were not detected from these noncoding sequence changes by proteomics, we believe that changes in enzymatic activity or DNA binding, in the case of regulators such as RelA and RpoS could provide incremental fitness contributions to tolerance in acetate. Indeed, a hierarchical clustering of expressed transcription factors (**Figure 5C**) corroborated a clear phenotypic grouping of the Σ-class strains that coincides with the different profiles we identified in our phenotypic characterization.

The Σ3 and Σ4 strains formed a cluster that is separate from Σ1, Σ2, and Σ5, indicating that global regulatory changes may impact growth on acetate. For instance, the higher relative abundance of the DNA-binding response regulator PP_1635 in Σ3 and Σ4 suggest a relationship with the observed shorter lag phases in comparison to the remaining strains, as there is a strong phenotypic association of this regulator with acetate conditions in the fitness browser dataset and in previous reports (29, 46). Interestingly, in this analysis, as well as in the whole proteome PCA clusterization, PT stands out as an outgroup based on several proteins that show higher than average percent abundance in comparison to all Σ strains. Particularly, HupB (PP_2303), the ß subunit of the histone-like protein HU, is markedly more abundant in PT than in other samples. HU is a nucleoid-associated protein found in prokaryotes as homo- or heterodimers of ɑ and ß subunits. The prevalence of different HU species is dependent on the growth phase and nutritional status of cells, and expression of HupB has been associated with exponential-phase growth in *P. putida* in glucose and succinate (50). HU activity as a regulator has been shown to promote tolerance against diverse stresses in bacteria, including cold-shock and oxidative stresses (51, 52), In *E. coli* the HU regulon comprises 8% of the genome and regulates the SOS system and the responses to several environmental stresses, including acidity and osmotic stress (53). While further characterization of HU activity in *P. putida* would be necessary to establish causal relationships between this regulator and phenotypic effects it promotes, its higher relative abundance in the PT background suggests that activation of different survival mechanisms in this strain might be a factor affecting growth performance of PT in acetate.

As differential protein expression provided more direct evidence for cellular effectors than the identifications mutations in global transcriptional regulators from the DNA resequencing dataset, we generated deletion strains in representative Σ-class strains Σ4 and Σ5 for candidate proteins that were upregulated in both strains, as we reasoned their increased expression was required for growth, and investigated how their absence affected fitness in glucose-containing media. We deleted *hemN* (PP_4264) and *gcvP-*I (PP_0988) in Σ4 and Σ5, and PP_2037 in Σ5. PP_2037 was not differentially expressed in the Σ4 background and excluded from testing. The *gcvP-I* and PP_2037 deletion mutants did not detectably impair growth in any of the conditions tested (**Supplemental Figure S5**), but the *hemN* deletion mutant resulted in a fitness defect in both strains with varied severity (**Figures 5D, 5E**). The *hemN* gene encodes an oxygen-independent coproporphyrinogen III dehydrogenase that plays a role in the biosynthesis of heme, an important cofactor for many protein functions including electron transport, oxygen transport, and relieving oxidative stress (54). In the Σ5 background, the *hemN* deletion mutant showed impaired growth on acetate alone, increasing lag phase, while when grown on the acetate-glucose mixed carbon source, this mutant was completely inviable for the duration of the kinetic growth assay. The Σ4 Δ*hemN* mutant showed at most a subtle growth defect in the same media conditions compared to the parental Σ4 strain. As the Σ4 background has many more proteins that were differentially expressed in the acetate medium, it is possible that different proteins could contribute to tolerance in a redundant manner, masking the Δ*hemN* phenotype in that background. This is also consistent with the divergent regulatory protein expression profiles where Σ4 and Σ5 fall into separate clusters (**Figure 5C**). We conclude at minimum from this analysis that HemN activity can enhance growth under acetate and isoprenol stress, and identify *hemN* as target of interest to understand the potential relationship of this oxidase with growth under acetate medium.

## Discussion

This is the first study that utilizes *P. putida* for the production of isoprenol using a defined glucose-acetate mixed carbon source modeled after substrate ratios (30, 31) found in lignocellulosic carbon streams. On its own, acetate is a poor target for producing biofuels since 3 molecules of acetate (a two-carbon molecule) would be necessary to generate a single molecule of isoprenol (a five-carbon molecule), requiring energy intensive steps in carbon-chain elongation. However, as a co-substrate abundant in lignocellulosic hydrolysate where glucose is readily generated, *P. putida* can use acetate to support biomass formation without compromising isoprenol titers. This context justifies the utilization of acetate as an ready co-substrate in an eventual biomanufacturing process as acetate and other organic acids are sometimes isolated from emerging “lignin first” deconstruction processes (55).

The main outcome of this study is that the acetate tolerant mutants fell into two phenotypic classes, as differentiated against their shortened lag phase, isoprenol productivity, and proteomics-based clustering. Acetate tolerance is a pleiotropic phenotype with many possible evolutionary routes, as seen with the mutants recovered here as well as in other bacteria. In *E. coli,* a different class of spontaneous acetate suppressor mutants were recovered with ALE for use in ethanol production (56). Other general approaches using barcoded mutant profiling (57) and saturation mutagenesis screening targeting the carbon catabolite repressor *crp* (58) have identified other acetate suppressor mutants. Towards our goal of acetate to isoprenol bioconversion in *P, putida*, the enhanced role of HemN in the Σ5 strain and its modest impact in Σ4 strain (refer to **Figure 5D**, **5E**) further reinforces the variable role cellular effectors can play. Other mutations, like a P387L substitution in *fleQ* (PP_4373) could be co-selected as a consequence of the laboratory growth conditions where strains are cultured with continuous shaking in aerobic conditions, leading to the accumulation of mutations in energy-intensive biological processes (59) which have previously been shown to impart metabolic costs in bioproduction strains (60). In the context of interconnected regulatory networks, identifying specific interventions that contribute to tolerance/production can be challenging, and ultimately Σ4 was not among the best isoprenol producers in mixed carbon source medium, despite being a fast grower (**Figure 2**, **Figure 3**). Still, the ease of generating suppressor mutants and their downstream analysis by proteomics justifies the bespoke nature of screening for acetate tolerized clones in specific hosts engineered to produce new biomolecules.

Importantly, our results suggest that acetate tolerization did not come at the cost of reduced isoprenol productivity. This specific substrate/product pair could have shown tradeoffs in isoprenol productivity as alcohol and acetate catabolism are reportedly linked in cellular metabolism in both *P. aeruginosa* (61) as well as *P. putida* (29, 62). In fact, while few proteins were found to be differentially expressed between PT and WT in acetate as a single carbon source, we found the alcohol dehydrogenase encoded by PP_2674 (PedE) to be significantly upregulated in PT in comparison to WT (log_2_FC = 2.6775; p-adjusted 0.03). PedE is one of two pyrroloquinoline quinone-dependent alcohol dehydrogenases that act on the oxidation of a variety of volatile organic compounds (VOCs) in the cell periplasm (63) in *P. putida*. A direct role of PedE in acetate catabolism seems unlikely, but its upregulation suggests a connection between acetate and alcohol metabolism, which would be consistent with ΔPP_2675 mutation impairing growth on acetate.

Determining the final cultivation parameters is an important consideration for any bioconversion process. In *E. coli*, bioconversion of glucose as a sole carbon source to isoprenol leads to accumulation of acetate as an overflow metabolite (21). This accumulation decreases the overall isoprenol yield where acetate forms an irreversible “dead end” product (21, 64). If glucose and acetate are both supplied in the medium as substrates, glucose is preferentially consumed first until depleted prior to acetate catabolism (65). In contrast, *P. putida* does not secrete acetate as an overflow metabolite (33, 66) and displays simultaneous consumption of both carbon sources under minimal salt medium growth conditions (**Figure 1**, **Figure 2**). Leveraging our mutant Σ-class strains, *P. putida* provides an advantaged microbial chassis for such substrates. As xylose- and galactose-enabled strains have also been established in *P. putida* KT2440 by several groups (67–70), future works could evaluate if specific metabolic engineering strategies are more efficient over the others towards the goal of complete carbohydrate-derived and aromatic-derived carbon streams (*ie,* unifying catabolic pathways and strain background optimization (67–70) to maximize isoprenol production from all available carbon in renewable biomass.

## Data availability statement

The *P. putida* sequencing data is available in NCBI SRA under the Bioproject Accession ID PRJNA1153078. The generated mass spectrometry proteomics data have been deposited to the ProteomeXchange Consortium via the PRIDE partner repository (71) with the dataset identifier PXD055153. [Information for the reviewer to login and review this information has been provided in the cover letter of this manuscript.] All other primary data is available from the corresponding author upon reasonable request. Plasmid sequences are available via the public JBEI registry, accessible at https://public-registry.jbei.org/ upon making a free login account.

## Funding Information

GMVS and M-EG were supported by the São Paulo Research Foundation (FAPESP) grants 2019/25432-7, 2021/01748-5 and 2022/12050-1. GMVS, TE, AM, TSL, YC, JJG acknowledge funding and support by the Joint BioEnergy Institute, U.S. Department of Energy, Office of Science, Biological and Environmental Research Program under Award Number DE-AC02-05CH11231 with Lawrence Berkeley National Laboratory.

## Acknowledgements

We thank all members of the Mukhopadhyay group and the Biofuels and Bioproducts Division at JBEI for their constructive comments on this project. We acknowledge technical assistance for chromatography analytical methods from Dr. David Carruthers (JBEI/LBNL).

## CRediT Statement

Conceptualization of project: TE, GMVS, AM. Experimental design, execution, and interpretation of data: GMVS, TE, AM, AS. Gas Chromatography data processing: AS, GS. Proteomics Data Pipeline: JJG, YC, CJP. Preparation of Figures: GMVS. Initial Draft of Manuscript: GMVS. Editing Manuscript: TE, GMVS, AM. Contributed Reagents: TE, TSL. Project Supervision: TE, AM. Raised Funds: M-EG, AM. All authors have read, given feedback, and approved the manuscript for publication.

## Notes

### Competing Interest Statement

The authors have declared no competing interest.

