## Supplementary material for "Alternate routes to acetate tolerance lead to varied isoprenol production from mixed carbon sources in *Pseudomonas putida*"

### Tables

**Table S1.** Summary information about the isolation conditions of the recovered Σ-class strains……1

**Table S2.** Maximum isoprenol titers (mM) detected for Σ-class and PT strains grown in different growth media……4

### Figures

**Table S1.** Summary information about the isolation conditions of the recovered Σ-class strains

| Strain | Experimental Batch # | Initial concentration of isoprenol in media | Validated Growth in M9 Isoprenol 11 g/L? | Notes |
|---|---|---|---|---|
| Σ1 | 2 | 8 g/L | Yes | - |
| Σ2 | 4 | 9 g/L | Yes | - |
| Σ3 | 5 | 8 g/L | Yes | Tolerized with pTE452 (pORTMAGE-Pa1) |
| Σ4 | 5 | 7 g/L | Yes | Tolerized with pTE452 (pORTMAGE-Pa1) |
| Σ5 | 5 | 9 g/L | Yes | Tolerized with pTE452 (pORTMAGE-Pa1) |

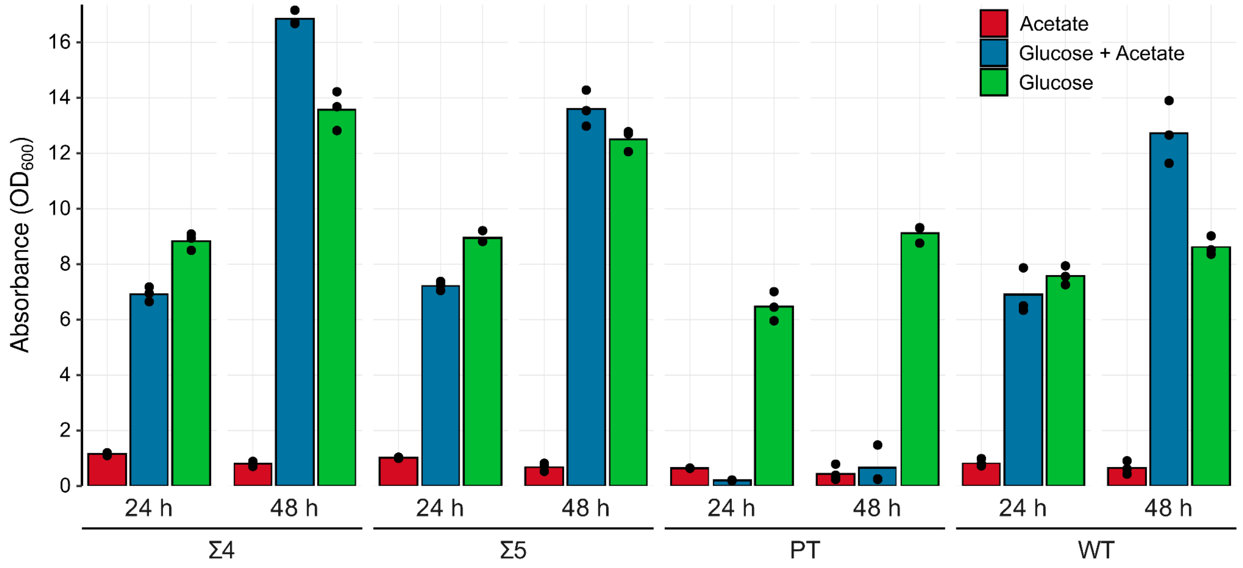

**Figure S1. Cell densities reached at 24 and 48 hours of cultivation for representative strains Σ4 and Σ5, WT and PT grown under different carbon sources.** Cultures were carried in test tubes using minimal media supplemented with acetate 4% (w/v), model hydrolysate medium composed of glucose 2% (w/v) and acetate 0.65% (w/v), or glucose 2% (w/v). Colored bars represent the mean OD value for three biological replicates in each condition. The absorbance values for each individual replicate are represented by the black points on the plot. Substrate consumption data for these cultures is plotted in **Figure 1** in the main text.

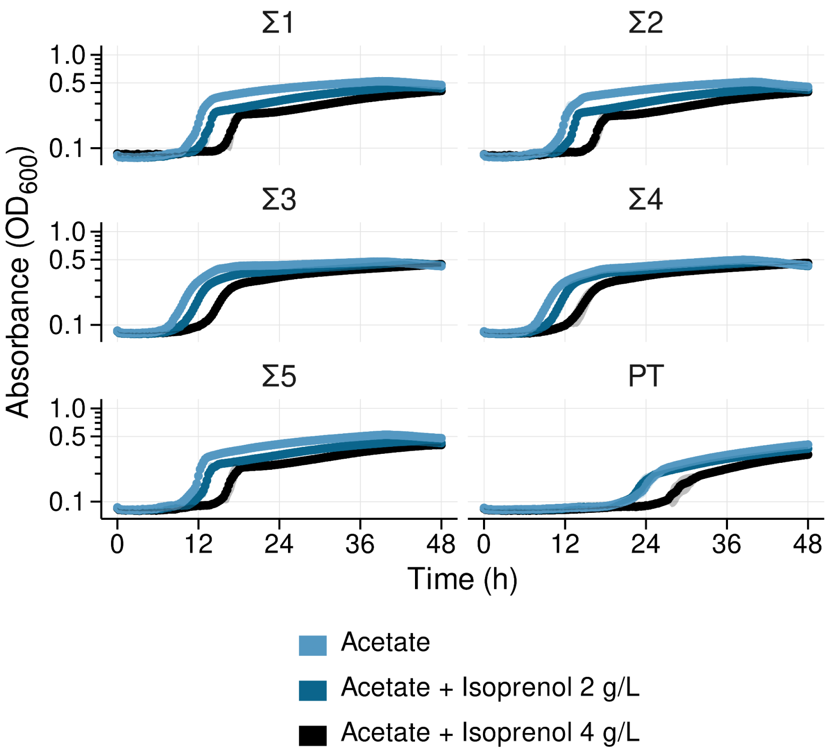

**Figure S2. Growth characterization of *P. putida* PT and Σ-class strains in M9 Acetate supplemented with isoprenol.** Panels display kinetic growth curves of cells cultivated in the microtiter dish format. All Σ strains display improved fitness in comparison to PT even when Isoprenol 4 g/L is supplemented to the medium. Despite this, growth of Σ strains is affected by the supplementation of as little as 2 g/L isoprenol in the medium, in comparison to the acetate alone condition. The color-shaded areas around the line represent the standard deviation of at least three biological replicates.

**Table S2.** Maximum isoprenol titers (mM) detected for Σ-class and PT strains grown in different growth media. In each column, the total amount of carbon ("C") in the medium, calculated as the sum of concentrations of all available carbon sources, is expressed in parentheses. n.d. = not determined. Glucose was always used at a concentration of 111 mM.

| | Medium | | | |
|---|---|---|---|---|
| | **M9 Acetate (75 mM "C")** | **M9 Acetate (100 mM "C")** | **M9 Glucose (111mM "C")** | **M9 Glucose + Acetate (221 mM "C")** |
| KT2440 "PT" | 0 | 0 | 3.29 | 0.05 |
| Σ1 | 0 | 0 | 4.52 | 1.44 |
| Σ2 | n.d. | n.d. | 4.51 | 1.34 |
| Σ3 | 0 | 0 | 2.41 | 1.05 |
| Σ4 | n.d. | n.d. | 2.39 | 1.23 |
| Σ5 | n.d. | n.d. | 4.26 | 1.88 |

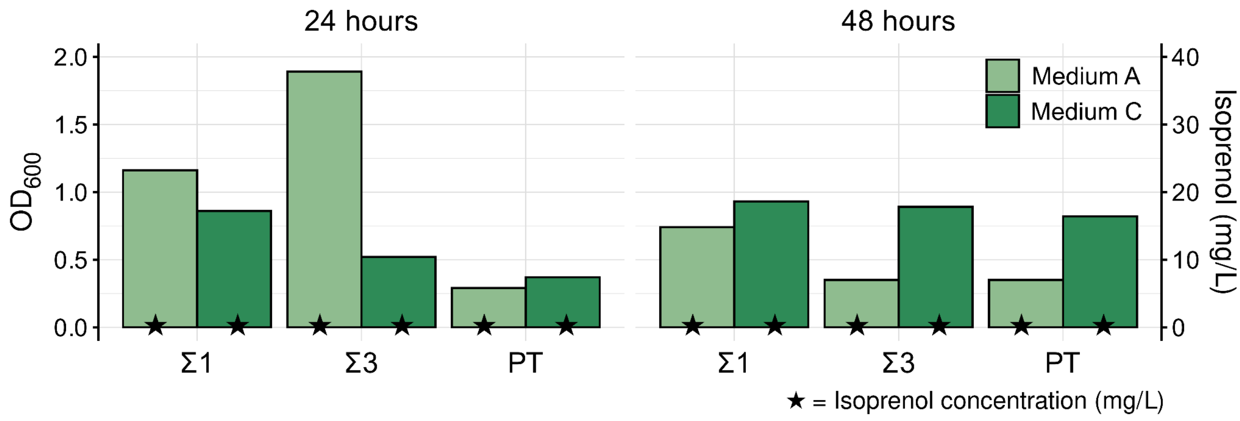

**Figure S3. Acetate supports growth, but not isoprenol production in Σ-class strains.** Heterologous isoprenol production in strains Σ1, Σ3 and PT in M9 acetate media (**Material and Methods**). Cells were grown for six hours before induction with arabinose 0.2% (w/v). Samples were harvested 24 and 48 hours after induction to monitor biomass formation ($OD_{600}$, green bars, left axis) and isoprenol concentration (black stars, right axis). For each strain, two M9 media compositions using varied acetate and ammonium sulfate concentrations. Medium A contained 75mM acetate and 75.5 mM ammonium sulfate. Medium C contained 100mM acetate and 1mM of ammonium sulfate. We were not able to detect isoprenol production by any of the strains under these conditions.

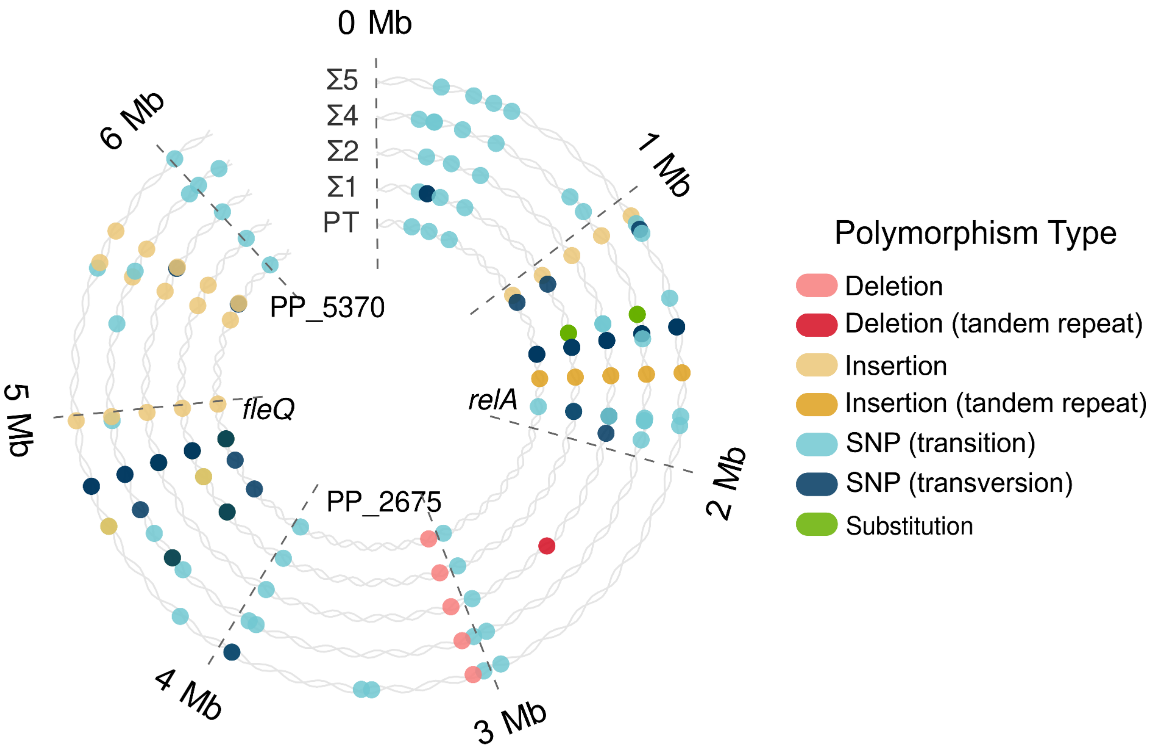

**Figure S4. An overview of mutations present in coding and non-coding regions of the genomes of Σ-class strains and PT in comparison to WT.** Several lineage-specific single nucleotide polymorphisms (SNPs) were identified that could contribute to acetate tolerance. Two background mutations preexisting in the PT strain but absent in WT P. putida KT2440 included a (+GG) insertion upstream of PP_4387 and a tandem repeat in PP_1458, affecting codon T180. These suppressor mutations may compensate for gene activity needed for the acetate response, though the role of these encoded proteins in metabolism remains unknown. The Σ strains showed various SNPs, such as A60T in PP_1623/rpoS for Σ5 and L312P in PP_1649 (D-lactate dehydrogenase) for Σ4. RpoS inactivation is linked to increased acetate fitness (Price et al., 2018), while PP_1649 transposon mutants show no fitness impact. Σ4 also harbored a Y399H mutation in PP_0184/argH, which was also linked to acetate fitness defects. Strains Σ4 and Σ5 had more coding mutations, likely due to the expression of mutL we included to increase the mutation frequency in these strains. PT carried a G203D mutation in PP_1656/RelA, while Σ1, Σ2, and Σ4 independently acquired further mutations in this gene. A comprehensive list of all mutations detected can be found in **Supplemental File 2**.

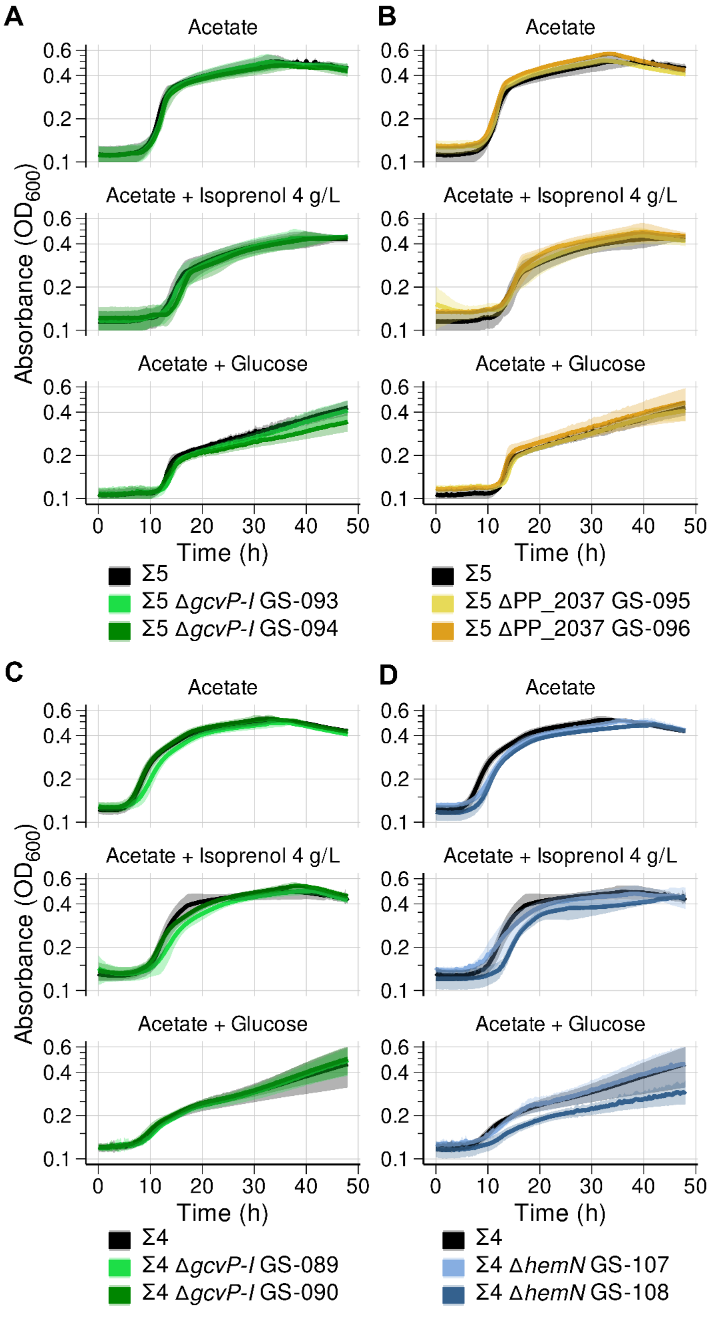

**Figure S5. Microplate-based growth kinetics of strains with deletions for *hemN*, *gcvP-I* and PP_2037 in Σ4 and Σ5 backgrounds.** In the panels, the darker lines represent the mean and the shaded areas the standard deviation of at least three independent biological replicates. For each deletion, two clones were screened for assessing true phenotypical differences caused by the absence of each gene. Σ4 Δ*hemN* (GS-108) data shown here WAS replotted against the parent and other isolated deletion clone generated via recombineering for the ease of comparison.
